# Blood-Mediated Opto-Acoustic Stimulation of Brain Cortex at Sub-millimeter Precision

**DOI:** 10.64898/2026.08.06.743316

**Authors:** Guo Chen, Mingsheng Li, Martin Thunemann, Kivilcim Kilic, Xinrui Gong, Carolyn Marar, Nan Zheng, Dingcheng Sun, Yueming Li, Fukai Chen, Hongru Zeng, Ji-Xin Cheng, Chen Yang

**Affiliations:** Department of Electrical and Computer Engineering, Boston University, Boston, MA 02215, USA; Department of Biomedical Engineering, Boston University, Boston, MA 02215, USA; Division of Materials Science and Engineering, Boston University, Boston, MA 02215, USA; Department of Biology, Boston University, Boston, MA 02215, USA; Department of Chemistry, Boston University, Boston, MA 02215, USA

## Abstract

Direct modulation of neural activity with high spatiotemporal precision is a cornerstone in experimental neuroscience. Here, we present a blood-mediated optoacoustic stimulation (BOAS) approach that utilizes blood as an endogenous transducer for brain stimulation. By delivering 532-nm nanosecond pulsed laser to the cortex, we demonstrate that the absorption of hemoglobin generates sufficient acoustic pressure to trigger neuronal activity. By integrating BOAS with calcium imaging in *GCaMP6f*-expressing mice, localized neuronal responses were observed. Quantitative analysis reveals that BOAS produces responses comparable to natural visual stimulation and is significantly more efficient than the photothermal stimulation. Furthermore, we show that the response is dose-dependent. At high energy doses, BOAS induces cortical spreading depression. Histological evaluation confirmed that the brain maintains tissue integrity even under these stimulation parameters. Together, this work establishes a versatile method for precise brain stimulation as an alternative method for stimulating neuron at cortex.

## Introduction

Coupling between changes in neural activation and hemodynamic signals has been a powerful tool for detecting neuronal activation in the brain^1^. Various imaging methods use the relationship between neuronal firing and hemodynamics to map brain activity. For example, functional magnetic resonance imaging (fMRI) maps neuronal activation by measuring the blood-oxygen-level-dependent (BOLD) signal, a contrast driven by the shifting ratio between diamagnetic oxyhemoglobin and paramagnetic deoxyhemoglobin in the venous bed^2^. Complementing fMRI, functional near-infrared spectroscopy (fNIRS) optically probes the same oxygenation shift by quantifying concentration changes in de-/oxyhemoglobin through their distinct absorption spectra, offering a highly portable, motion-tolerant tool for superficial cortical mapping in naturalistic settings^3^. More recently, functional ultrasound (fUS) imaging has emerged as a powerful modality for mapping microvascular cerebral blood volume (CBV) dynamics. It captures the hemodynamic correlates of neural activity with exceptional spatiotemporal sensitivity for high-resolution, deep-brain imaging in both awake preclinical models and intraoperative human applications^4^.

In addition, photoacoustic (PA, also called optoacoustic) imaging has been developed as a functional imaging technique with high efficiency and non-invasive nature^5^. The photoacoustic process converts absorbed optical energy into ultrasonic waves. It has enabled a broad range of applications across photoacoustic microscopy (PAM)^5-9^ and tomography (PAT)^9-11^, offering high-contrast visualization of vasculature, oxygenation, and metabolic activity *in vivo*. Because hemoglobin in the blood is a strong optical absorber whose spectrum depends on oxygenation state, multispectral PA imaging can map functional parameters such as blood oxygen saturation (sO_2_) and hemodynamic changes with high spatiotemporal fidelity^12,13^. These structural and functional capabilities have made PA imaging an increasingly powerful tool in neuroimaging. By visualizing cerebral vasculature, local blood-volume and sO_2_ changes, as well as fast hemodynamic responses that correlate with neural activity, PA imaging links optical contrast to neurophysiology across microscopic to macroscopic scales^9^. However, whether one could utilize PA effects to enable functional control over neural activity is yet to be investigated.

Here, we present a blood-mediated optoacoustic stimulation (BOAS) approach, representing the first demonstration of using endogenous absorbers as a photoacoustic emitter for neural stimulation. We first demonstrate that the PA pressure generated by blood is comparable to that of previously developed carbon nanotube/polydimethylsiloxane films, an efficient PA emitter reported for robust neuromodulation^14^. This comparison laid a solid foundation for using blood as an endogenous PA stimulator. Then, we developed an integrated system capable of delivering nanosecond-pulsed laser excitation to induce PA generation in cortical blood vessels while simultaneously performing widefield calcium imaging for monitoring neuronal activity in the cortex using the genetically-encoded calcium indicator GCaMP6f. This configuration enabled real-time stimulation and monitoring of neural activities in the rodent brain. We observed robust calcium responses in the visual cortex following BOAS, confirming effective neural activation. Importantly, compared with purely photothermal stimulation conditions under the same laser energy, BOAS elicited stronger neural activation, suggesting that the optoacoustic mechanism enhances stimulation efficiency, lowering the stimulation threshold and reducing thermal toxicity. This new technology opens up new potentials for highly efficient non-genetic modulation of brain activity.

## Results

### Blood generates sufficient acoustic pressure for neural stimulation

The high optical absorption coefficient of hemoglobin has firmly established blood as an exceptionally efficient photoacoustic contrast agent for high-resolution functional imaging^11^. Here, we evaluated whether sufficient acoustic pressure can be generated by hemoglobin to use blood as an endogenous PA stimulator to evoke activity of adjacent neurons (**Figure 1a)**. We performed the initial PA performance characterization in whole bovine blood (BOVINE BLOOD DEFIB 100ML, Hemostat Laboratories) and compared it to that of a carbon nanotube/polydimethylsiloxane (CNT/PDMS) thin film under the same laser condition. The CNT/PDMS thin film sample was selected as it is a well-characterized, high-efficiency photoacoustic emitter previously validated for exogenous neuromodulation *in vitro* and *in vivo*^14,15^. The excitation wavelength of 532 nm is chosen because it is an isosbestic wavelength for oxyhemoglobin and deoxyhemoglobin^16^, allowing effective excitation to blood in veins and arteries. The extinction spectrum of whole bovine blood measured with a spectrometer is shown in **Figure S1**. As shown in **Figure 1b**, using a 532-nm, 0.5-ns pulsed laser (850 nJ/pulse, 20 kHz), the bovine blood sample produced a PA signal with an amplitude of 112 mV measured by a 25-MHz ultrasound transducer (V324-SM, Olympus), compared with 142 mV for the CNT/PDMS emitter. Although the PA amplitude from blood is lower, the difference is modest, indicating that blood is a promising endogenous absorber capable of generating sufficient PA pressure for stimulation.

**Figure 1.**
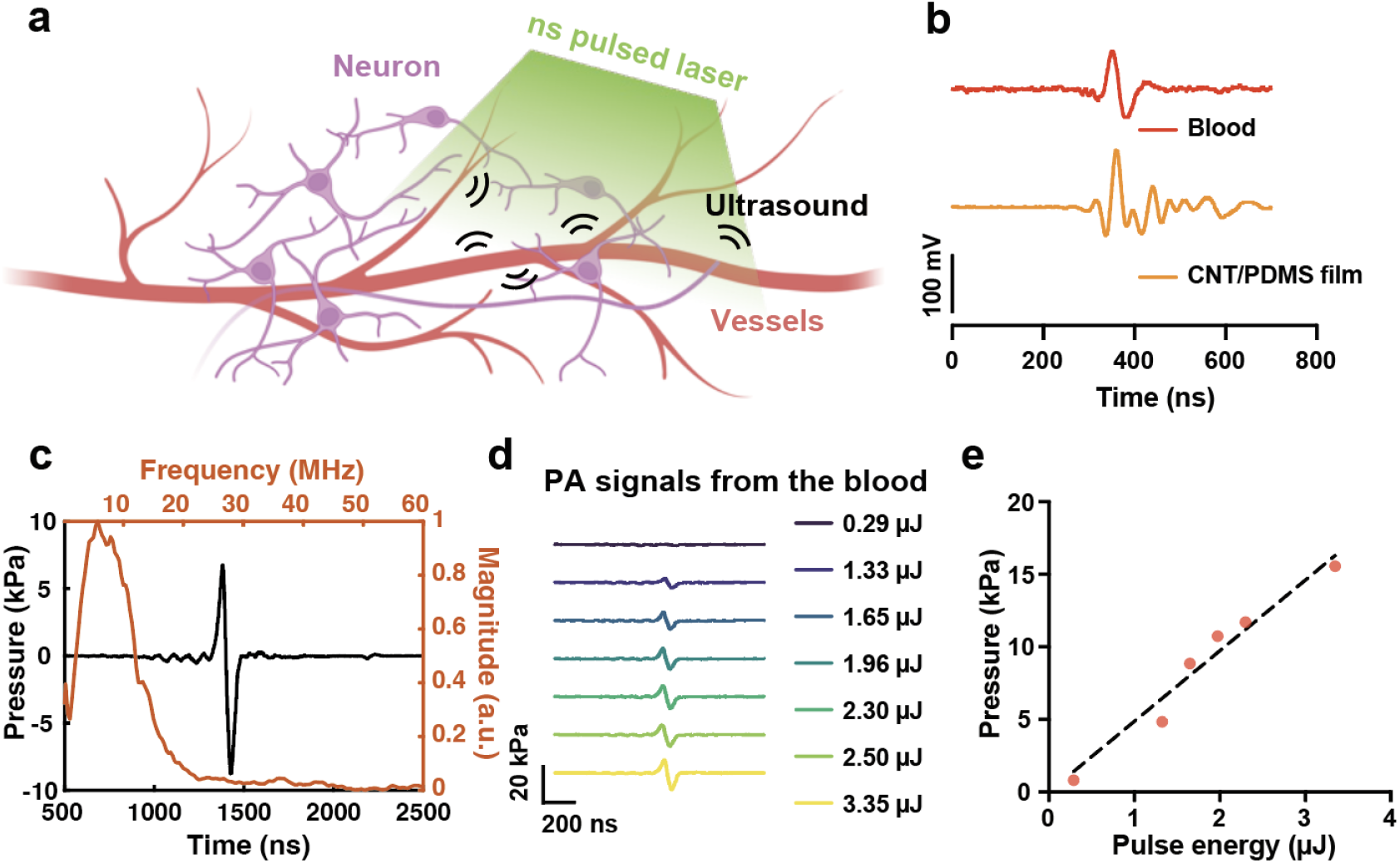
Blood as an endogenous absorber for photoacoustic (PA) wave generation. **a**. Schematic showing blood as an endogenous absorber for PA-mediated neuronal activation. **b**. Comparison of the PA signal generated by a bovine blood sample and a CNT/PDMS film (single pulse, 850 nJ). **c**. The PA signal from a bovine blood sample plotted as a function of time (black) with frequency spectrum (after FFT) of the PA signal in red. Laser condition: 532 nm, 3.35 μJ. **d**. PA signal generated by a whole blood sample with different pulse energies. **e**. Peak-to-Peak PA pressure as a function of laser pulse energy.

To further quantify the PA pressure generated by blood, we measured the photoacoustic signal of whole bovine blood using a calibrated hydrophone (HNR1000, Onda). Under laser excitation at 532 nm with a 0.5-ns pulse width and 3.35 μJ pulse energy, blood produced a peak-to-peak pressure of 15.5 kPa with a central frequency of approximately 5 MHz (**Figure 1c**). When the incident laser energy was varied, the resulting PA pressure scaled proportionally, showing a linear dependence with a slope of 4.8 kPa/μJ (**Figure 1d–e**). These results provide critical evidence that hemoglobin-generated PA pressure reaches the necessary threshold for mechanical membrane depolarization in nearby neurons.

### An integrated system for cortical BOAS and calcium imaging

We developed an integrated optical platform capable of simultaneous stimulation using blood as an endogenous absorber and real-time monitoring of neuronal activity in mouse visual cortex using calcium imaging as a robust imaging method that has previously been used to observe ultrasound-induced changes in neuronal activity^17^.

As illustrated in **Figure 2a**, we designed a setup that combines a custom epi-fluorescence imaging system with excitation at 488 nm with a 532-nm nanosecond pulsed laser (0.5 ns pulse width, 20 kHz repetition rate) for PA generation in blood. The excitation light for calcium imaging and the pulsed laser for stimulation are co-aligned and delivered through a single 4x objective lens, and weakly focused onto the mouse cortex through a cranial window. Specifically, we integrated a high-rejection notch filter (NF) into the detection path to effectively eliminate laser-induced artifacts detected by the camera, ensuring high-fidelity recording of calcium dynamics while the blood-mediated PA stimulation is active. This integrated approach allows for the direct correlation of endogenous PA modulation with immediate neuronal calcium responses across the cortical surface.

**Figure 2.**
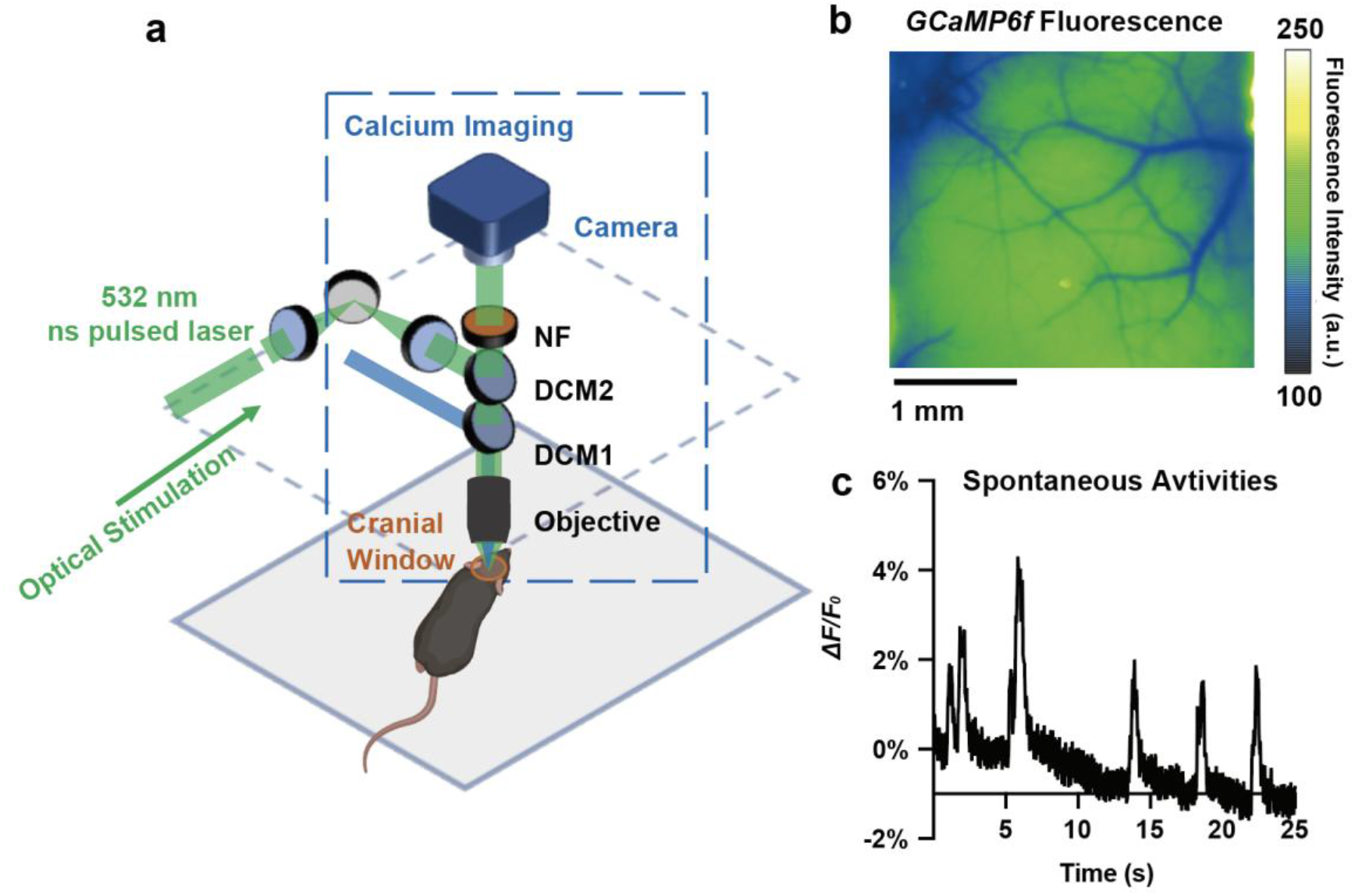
An integrated system for cortical optical stimulation and calcium imaging. a. Schematic of the system. Blue box: calcium imaging. NF: notch filter. DCM: dichroic mirror. b. Representative calcium imaging of the mouse cortex. Dark area: vessels. Scale bar: 1 mm. c. Spontaneous calcium activity under anesthesia.

To monitor the brain activities in a live animal, we performed a chronic cranial window implantation over the visual cortex of wild-type mice after local delivery of adeno-associated virus to induce neuronal expression of *GCaMP6f* (details are described in the Method section). This surgical preparation provided an optically clear interface to the brain, allowing for the simultaneous visualization of the cortical vasculature and fluorescent signals from *GCaMP6f* resembling neural dynamics within the visual cortex. A representative wide-field fluorescence image of the cortex through the cranial window is shown in **Figure 2b**. Blood vessels appear dark against the bright fluorescence background, as hemoglobin within the vessels absorbs the emitted fluorescence signal. To validate the performance of our customized imaging system, we recorded calcium activity in the mouse brain under anesthesia. As shown in **Figure 2c**, the system successfully captured robust spontaneous neuronal activity. This confirms that the integrated setup maintains high sensitivity and temporal resolution for monitoring neural dynamics *in vivo*.

### BOAS successfully induces changes in neuronal activity in mouse visual cortex

To evaluate the feasibility of blood-mediated optoacoustic modulation of neuronal activity *in vivo*, we designed an experimental protocol to compare our photoacoustic (PA) approach with a sensory stimulation. As detailed in the experimental setup in **Figure 3a**, a 532-nm pulsed laser was weakly focused onto the cortical surface, creating a localized stimulation site with a spot diameter of approximately 375 µm. The laser profile is shown in **Figure S2**. The ns-pulse laser was operated at a power of 200 mW with a pulse train duration of 100 ms. As a positive control, we delivered a visual stimulus consisting of a blinking white-light LED (2 Hz, 2-s duration) to the contralateral eye of the mouse. We monitored and compared the resulting neuronal calcium dynamics in the visual cortex in response to both PA and visual stimulation for a direct assessment of the efficacy and spatial localization of BOAS relative to natural sensory processing.

**Figure 3.**
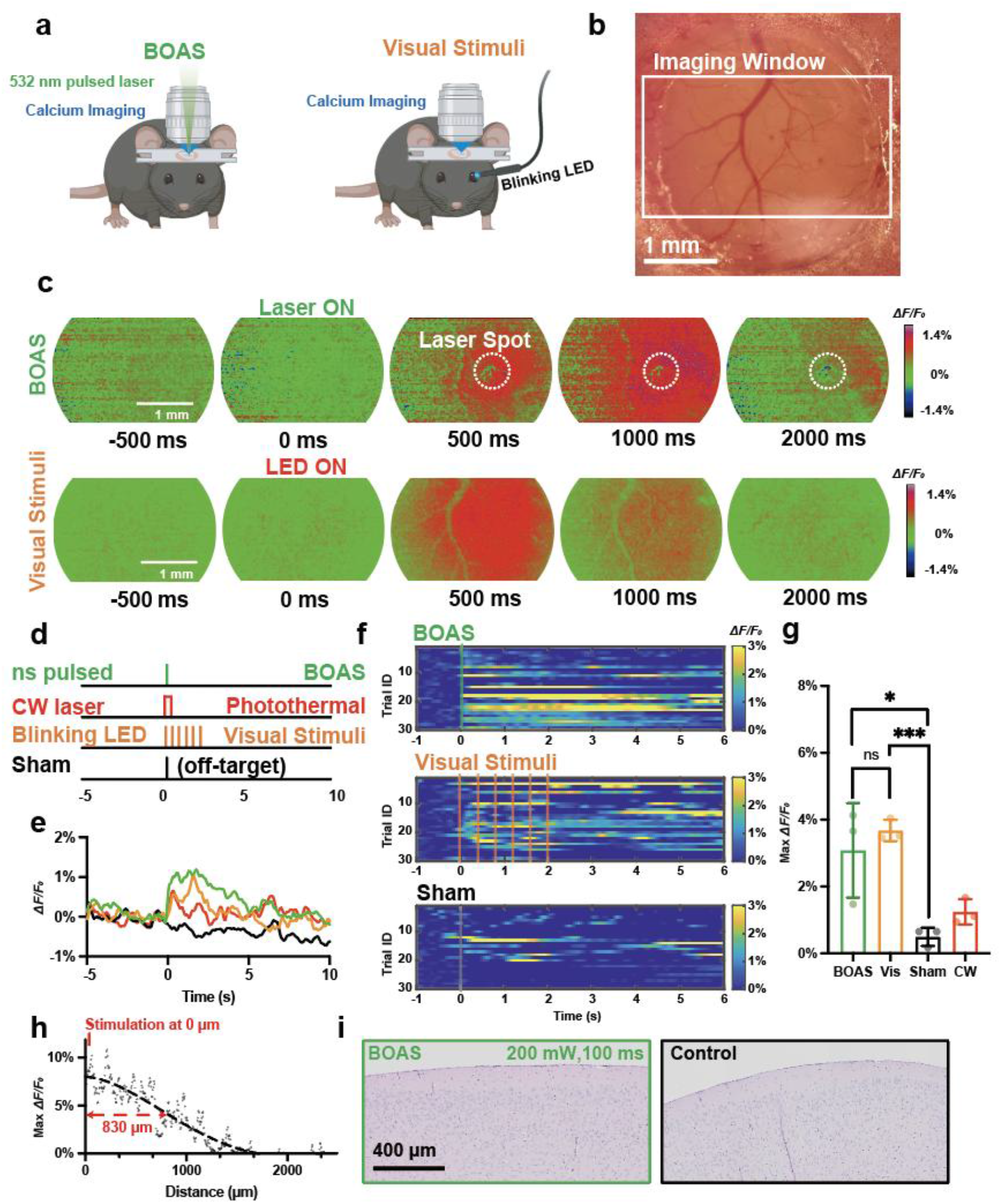
Blood optoacoustic stimulation (BOAS) of mouse brain cortex monitored by calcium imaging. **a**. schematic of BOAS and visual stimuli as a positive control. **b**. cranial window above mouse visual cortex for imaging and stimulation (centered at 2.6 mm posterior, 2 mm lateral to Bregma). **c**. representative image of fluorescence intensity changes upon stimulation. N=10 trials for BOAS and visual stimulation. BOAS: 532 nm, 200 mW, 0.5 ns pulse width, 20 kHz repetition rate, 100 ms burst duration. Visual stimuli: white light, 100 ms, 3 Hz, 2 s. **d**. PA stimulation, PT stimulation, positive control, and Sham control protocol. **e**. averaged calcium trace of BOAS and visual stimuli. N=30 from 3 animals for both conditions. BOAS: 532 nm, 200 mW, 0.5 ns pulse width, 20 kHz repetition rate, 100 ms burst duration. Visual stimuli: white light, 100 ms, 3 Hz, 2 s. **f**. trial by trial traces of BOAS, visual stimuli, and sham control. Dash line indicates when the laser / LED was turned on. **g**. Statistical analysis between groups. N=3 animals for each condition. n.s.: not significantly different. *: p<0.05. ***: p<0.001.. **h**. Maximum calcium response plotted as a function of distance away from the stimulation site. **i**. Histology results for mice with BOAS treatment (200 mW, 100 ms) and without treatment.

The cranial window image is shown in **Figure 3b**. The resulting cortical calcium dynamics are presented in **Figure 3c**, comparing photoacoustic (PA) stimulation (top row) with visual/sensory stimulation (bottom row). After the 532-nm laser was turned on at t = 0 s, a calcium transient was observed in the BOAS condition within 0.5 s. The evoked neuronal activity was spatially localized, concentrated in the cortical area surrounding the laser spot. As plotted in **Figure 3h**, the stimulated area is larger than the laser spot size, with a radius of 0.83 mm. This demonstrated that our BOAS has sub-millimeter stimulation precision. Similarly, the positive control exhibited a clear response in the visual cortex following stimulation onset at t = 0 s. Both stimuli successfully elicited changes in neuronal activity, as evidenced by the increase in *GCaMP6f* fluorescence intensity. These representative imaging results, averaged over 10 trials on the same animal, demonstrate that blood-mediated PA stimulation can achieve activation comparable to natural sensory pathways, confirming its efficacy as a localized neuromodulation tool.

To rigorously quantify the efficacy of BOAS, we compared calcium dynamics across multiple subjects and trials, as summarized in **Figure 3d-e** (BOAS was repeated in three different animals, and 10 repeated trials for each animal. The visual stimuli and sham control were done on the same three mice for 3×10 trials in total). The temporal profiles of fluorescence changes under different experimental conditions (**Figure 3e**) illustrate a clear divergence between the experimental and the sham control conditions, where an off-target laser treatment with the same power and duration was delivered (**Figure S3**). Both the BOAS condition (green) and the visual stimuli condition (orange) exhibited robust, synchronized calcium responses upon stimulation, while the sham control (black) showed no detectable change in activity. These data represent the mean response across 30 trials (10 trials per mouse, three mice in total). The corresponding temporal profiles of the stimulation protocols, including the laser pulse train and LED timing, are provided in **Figure 3d**.

In addition to the averaged temporal profiles shown in **Figure 3e**, the consistency of the evoked responses across all experimental repetitions is illustrated in **Figure 3f**. Here, the calcium traces for each individual trial from BOAS, visual stimulation, and sham control are presented as heatmaps, where the x-axis represents time and each row corresponds to a single trial (trial ID 1-10 is from mouse no.1, 11-20 is from mouse no. 2 and 21-30 is from mouse no.3). These heatmaps reveal a high degree of trial-to-trial reliability and temporal synchronization with the stimulus onset (t = 0 s), further supporting the robustness of blood-mediated photoacoustic neuromodulation.

Statistical analysis of the peak response (**Figure 3g**) further validates these findings. For the statistics, we averaged the 10 repeats in the same animal and calculated the statistical difference between each experiment condition with a biological repeat of 3 different mice. The BOAS reached a maximum *ΔF/F*_*0*_ of 3.08% ±1.42% , which was comparable to the 3.68%±0.32% observed in the visual stimulation conditions. A *t-test* revealed no significant difference between the BOAS and visual stimulation intensities (p=0.5184), suggesting that photoacoustic stimulation can elicit neuronal responses of a magnitude similar to natural sensory stimulation under anesthesia. Critically, both the BOAS and visual stimulation conditions produced significantly higher responses compared to the sham control (*: p=0.0359 and ***: p=0.0002, respectively), confirming that the observed calcium transients were specifically induced by the targeted stimulation.

To distinguish the contribution of the photoacoustic mechanical effect from pure photothermal heating, we conducted a control experiment substituting the nanosecond pulsed laser with a continuous-wave (CW) laser of the same wavelength (532 nm). By maintaining 100-ms exposure duration and 200 mW averaged power, we ensured that the total energy delivered was identical, while there will be only photothermal effect in the case of CW laser. As shown in **Figure 3e,f** (red), the CW photothermal condition elicited a detectable but markedly weaker neural response, reaching a maximum *ΔF/F*_*0*_ of only 1.24%±0.38%. The calcium imaging results and trial-by-trial traces can be seen in **Figure S4**. The amplitude of the response was lower than that achieved by the BOAS condition at the same energy by 59%. While these results suggest that photothermal effects alone are capable of inducing some modulation of neuronal activity, the addition of the photoacoustic mechanical component in the pulsed condition results in a significantly more efficient stimulation. This comparison underscores that acoustic waves generated from blood are the primary drivers of the robust neuronal activation observed in our system.

To evaluate the safety of the stimulation parameters used in this study, we performed histological analysis on cortical tissue following the experimental sessions. Hematoxylin and eosin (H&E) staining was conducted on brain sections harvested from PA-stimulated regions (200 mW, 100 ms pulse train). The selected area of the brain was sliced into 15 sections with 5 μm thickness, and all slices were examined to make sure no potential damage was missed. Representative images of the experimental and control conditions are shown in **Figure 3i**. The histological assessment revealed no detectable signs of tissue damage: compared to the control that received no laser treatment, the stimulated cortical tissue exhibited a similar morphology and structural integrity. The preserved cellular architecture and the absence of lesions in both conditions indicate that the endogenous photoacoustic transients were generated within a safe operating range, eliciting robust neuronal activity without compromising the underlying neural or vascular environment.

### Cortical response to BOAS under different laser energy dosages

To characterize the relationship between the total laser energy dosage delivered and the resulting neuronal activity, we evaluated cortical responses under varying laser durations. While maintaining a constant laser power of 200 mW, we modulated the total energy delivered by reducing the pulse train duration from 100 ms to 50 ms and 25 ms. As with previous experiments, each condition was rigorously tested across 30 repeated trials from three mice (10 trials on one mouse, three mice in total)

The individual trial dynamics are visualized as heatmaps in **Figure 4a**, categorized by duration: 50 ms (orange), and 25 ms (black). The corresponding averaged temporal traces are presented in **Figure 4b**, where the 100-ms condition shows an obvious stimulation result and the 25-ms condition shows a negligible response (the 100-ms data is adopted from **Figure 3**). Quantitative statistical analysis (**Figure 4c**) revealed a pronounced, duration-dependent decline in neuronal activation when the stimulation duration was shorter. Specifically, reducing the stimulation duration from 100 ms to 50 ms resulted in a substantial decrease in the calcium signal, with the peak *ΔF/F*_*0*_ dropping to 0.8%. At a duration of 25 ms, the peak signal further diminished to 0.3%, a value approaching the noise floor of the imaging system. These findings demonstrate that blood-mediated photoacoustic neuromodulation follows a clear dose-dependent response, allowing for tunable control over the intensity of induced brain activity.

**Figure 4.**
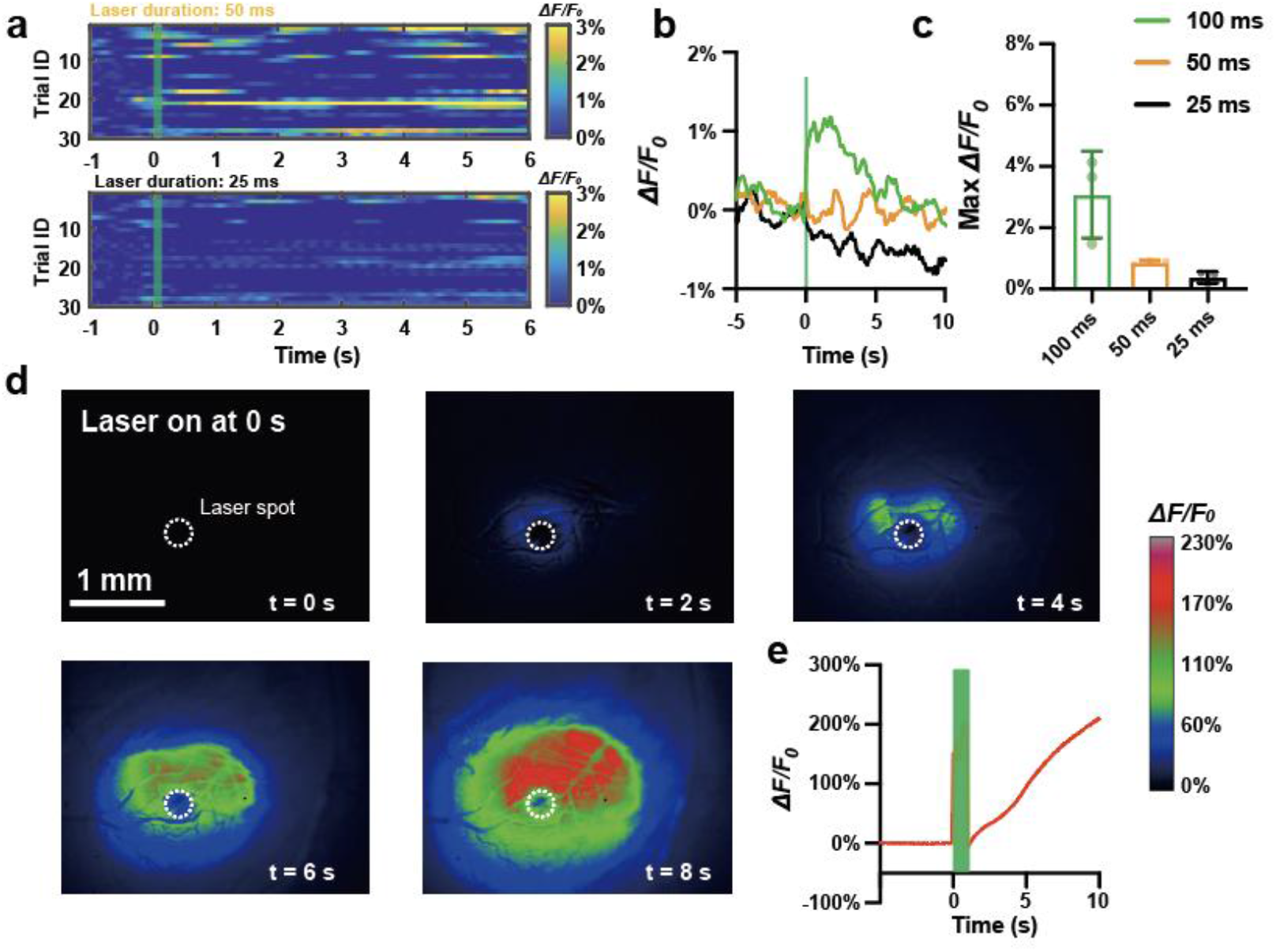
Cortical response to BOAS with different laser energy dosages. **a**. trial by trial traces of BOAS under 50 ms (orange) and 25 ms (black). **b**. calcium trace of the visual cortex plotted as a function of time. Green: 100 ms. Orange: 50 ms. Black: 25 ms. **c**. maximum fluorescence change of the visual cortex under different durations of BOAS. N=3 animals for each condition. **d**. Calcium imaging of the mouse’s visual cortex showing a cortical spread depression (CSD) under a laser power of 380 mW, and a duration of 1000 ms. **e**. Calcium trace of the visual cortex showing a cortical spread depression under the same laser condition as panel d.

Beyond localized neuromodulation, we investigated the cortical response to elevated laser energy dosages. Interestingly, when the laser power was increased to 380 mW with a prolonged pulse train duration of 1000 ms, the system elicited a qualitatively different neural phenomenon: as illustrated in **Figure 4d**, the stimulation triggered a pronounced calcium wave that originated at the laser targeting site and spread progressively across the visible cortex at an average speed of 5.6 mm/min. The spreading speed induced by BOAS is consistent with previously reported works showing Cortical Spreading Depression (CSD) propagation can vary from 1-8 mm/min^18^.

The temporal profile of this response, plotted as the red trace in **Figure 4e**, shows a high-amplitude calcium signal that persisted for a long time before gradually returning to baseline. This spatiotemporal pattern—characterized by a slow-moving, high-intensity wave of activity followed by a prolonged recovery period—is highly consistent with stimulation-induced CSD. These results indicate that while lower energy dosages provide precise, localized stimulation, blood-mediated photoacoustic effects at higher thresholds can be used to reliably initiate large-scale cortical events. Similar findings were reported in FUS (Focused Ultrasound) brain stimulation, where, under strong ultrasound stimuli, CSD was observed through calcium imaging of the mouse cortex ^19^. This result indicates that BOAS behaves similarly to other ultrasound-based techniques for brain stimulation.

## Discussion

With translational potential to modulate human brain activity, non-genetic brain stimulation remains a frontier of neurotechnology. Water in brain as an endogenous absorber has been utilized in infrared stimulation (INS) of neural activities in brain ^20^. By utilizing the water absorption peak in the near-infrared window, INS can directly heat up the water and generate a localized temperature increase, which modulates the brain through thermosensitive ion channels ^21,22^ or changing the membrane capacitance of neurons ^21,23-25^. Due to its non-genetic nature, INS has been applied on human patients for clinical treatments ^26^. However, the potential thermal toxicity caused by INS still raises concerns for its clinical trials, and new non-genetic methods are still sought out for neural stimulation^27^.

Photoacoustic neural stimulation through exogenous PA converters has been recently developed and applied to activate neurons in brain and retina. Jiang et al. demonstrated that a fiber-based optoacoustic emitter could successfully stimulate the mouse cortex ^28^. A multifunctional fiber-based emitter integrating an optical waveguide and recording electrodes has been implanted into the mouse hippocampus to provide a bidirectional interface capable of simultaneously stimulating and monitoring neural activity^29^. Tapered fiber–based PA emitters (TFOE) have further improved spatial precision, achieving single-cell–level stimulation and enabling tight correlation with electrophysiology using patch-clamp recordings in brain slices ^30^. Meanwhile, successful retinal modulation through PA has been demonstrated in vivo, showing the potential of non-genetic photoacoustic retina prosthesis^14^. Studies have indicated that photoacoustic stimulation can be more efficient than photothermal stimulation. Using the same fiber emitter for PA and PT generation, the PT condition needs 40 times larger total energy dosage to achieve similar stimulation conditions compared with PA stimulation, and the photothermal effect accompanying during the PA stimulation scenario is unable to open up thermal-sensitive ion channels nor significant capacitance changes of the membrane to induce action potentials^27^. Nevertheless, application of exogenous PA agents imposes risk of surgical procedure, immune response, and post-operation complications.

In this work, we introduce blood-mediated optoacoustic stimulation (coined “BOAS”) as a fundamentally new neuromodulation paradigm that leverages an endogenous light absorber, i.e. circulating blood, to generate PA pressure for brain stimulation. This approach overcomes a limitation of existing PA neuromodulation technologies, which require implanted or administered PA emitters that can require surgical procedures, impose risks and /or interfere with imaging modalities. By using blood itself as the PA source, BOAS provides a fully intrinsic, label-free stimulation mechanism that is naturally co-localized with the cerebrovascular network.

Our characterization experiments demonstrate that blood generates strong PA signals under ns-pulsed illumination. When benchmarked against a CNT/PDMS emitter, an existing PA neuromodulation device, blood produced only modestly lower PA amplitude, reinforcing its viability as a functional PA source for stimulation. Hydrophone measurements further revealed that blood-generated PA pressure scales linearly with laser energy and reaches peak pressures in the tens of kilopascals under conditions relevant to neuromodulation. These measurements establish the physical basis for BOAS and suggest that the blood as an endogenous absorber can readily support PA-based stimulation.

Using our integrated stimulation and imaging system, we could show in anesthetized mice that PA generation in cortical vessels reliably evoked neural activity *in vivo*. Importantly, the magnitude of calcium responses elicited by BOAS was comparable to those evoked by visual sensory stimulation and far exceeded sham conditions, confirming that the responses reflected genuine neural activation rather than imaging artifacts, hemodynamic confounds, or indirect stimulation of the animals’ visual circuit. BOAS also produced significantly stronger activation than continuous wave photothermal irradiation under equivalent energy dosages, suggesting that the mechanical pressure impulse characteristic of PA generation, rather than temperature alone, plays a critical role in neural excitation.

Beyond demonstrating efficacy, our findings highlight the tunability and dynamic range of BOAS. Decreasing laser duration reduced the PA-induced calcium response to near baseline levels, consistent with the reduced acoustic pulse at lower energy deposition. Conversely, high-dose stimulation produced widespread calcium elevation and a prolonged depression-like response consistent with CSD, illustrating the ability of BOAS to modulate neural activity across regimes—from localized excitation to large-scale network perturbations. This dose– response relationship offers a pathway toward flexible neuromodulation strategies tailored to specific experimental or therapeutic needs.

Importantly, the stimulation parameters used in our primary BOAS experiments did not induce detectable tissue damage, as supported by post-hoc histology, aligning with the safety profile expected of short-pulse optoacoustic excitation. Together, these results position BOAS as a promising and minimally invasive neuromodulation method that integrates seamlessly with optical neuroimaging.

Several limitations remain. First, although blood is ubiquitous in the brain, the spatial specificity of BOAS ultimately depends on the distribution of cortical microvasculature, which may constrain targeting resolution relative to fiber-based PA emitters or optogenetics. Second, because the current implementation is limited to superficial cortical regions accessible to optical excitation, adaptations for deeper targets may require alternative wavelengths, nonlinear optical strategies, or endoscopic delivery. Third, while our results strongly suggest a mechanistic contribution from the PA pressure wave, the exact biophysical pathway, for instance, whether via membrane mechanotransduction, ion channel modulation, or vascular– neuronal coupling, requires further investigation. Lastly, compared to optogenetics brain modulation, although BOAS avoids genetic modification, it also loses the cell-type specificity. Nonetheless, BOAS uniquely enables optoacoustic neuromodulation without introducing foreign implants or acoustic emitters, making it particularly advantageous for systems neuroscience studies requiring clean optical access and minimal tissue perturbation.

## Methods

### Fabrication of CNT/PDMS film

To fabricate the CNT/PDMS film for photoacoustic characterization, we followed the previously published study^14^. The PDMS was prepared at a mix ratio of 10:1 (base to curing agent). Then, a 15%wt of CNT (<8 nm OD, 2–5 nm ID, length 0.5–2 µm, VWR, Inc., USA) was mixed with the PDMS, with the addition of IPA to facilitate CNT dissolution. 5-minute sonication and 30-minute degassing steps were applied to remove the gas inside the mixture. The prepared mixture was then spin-coated onto a glass substrate at 500 rpm for 5 minutes to form a film of 40 µm. The coated substrate was cured at 110 °C for 15 minutes.

### Characterization of photoacoustic signals

The amplitude of the PA signal generated by bovine blood (BOVINE BLOOD DEFIB 100ML, Hemostat Laboratories) and the CNT-PDMS film were measured using a needle hydrophone (HNR1000, Onda) or a ultrasound transducer (V324-SM, Olympus). A digital oscilloscope (DSO6014A, Agilent Technologies, CA, USA) recorded the electrical signal from the hydrophone. A four-axis micro-manipulator (MC1000e controller with MX7600R motorized manipulator, Siskiyou Corporation, OR, USA), with a resolution of 0.2 μm, controlled the distance between the PA emitters and the hydrophone.

### Cranial window implantation and virus injection

All experimental procedures have complied with all relevant guidelines and ethical regulations for animal testing and research established and approved by the Institutional Animal Care and Use Committee of Boston University (PROTO201800534).

We used 3 male C57BL/6J (The Jackson Laboratories Animals) mice at an age of 14-16 weeks for the study. Animals were held in isolated ventilated cages either in groups or single-housed with unlimited access to food and water.

For cranial window implantation and AAV delivery, mice were injected with 4.8 mg/kg Dexamethasone 4 hours before surgery, and they further received Meloxicam and extended-release Buprenorphine for management of post-surgical pain and inflammation. For surgery, animals were anesthetized with isoflurane and placed on a feedback-controlled heating pad in a stereotaxic frame in prone position; a nose cone delivered 1-2% isoflurane in 100% oxygen for the duration of the surgery. After hair removal, skin was cleaned with 3 alternating applications of povidone-iodine and ethanol. A round piece of skin was removed to expose the skull; the retracted skin was attached to the bone using surgical-grade cyanoacrylate glue (VetBond, 3M). The skull surface was then treated with gel etchant containing 35% phosphoric acid (Kerr Dental) for 60-90 s, after which bone and surrounding skin were washed with sterile 0.9% NaCl. A custom-made titanium headpost was fixed to the bone using UV-curable dental resin (NX3, Kerr Dental) and composite (Tetrik EvoFlow, Ivoclar Vivadent). Then, a 3-mm round craniotomy was performed over the visual cortex (centered at 2.6 mm posterior, 2 mm lateral to Bregma) using a dental drill, and 150 nL AAV solution (AAV9.Syn.Flex.GCaMP6f.WPRE.SV40; Addgene, MA, USA) was delivered at 150-200 µm depth via tissue microinjection with a tapered glass pipette at 15 nL/min for 3 min (Nanoject III, Drummond). After injection, the glass pipette was left in place for 10 min to allow the AAV solution to diffuse into the tissue. Per animal, the AAV solution was injected into three different locations within the visual cortex for relatively uniform GCaMP6f expression within the exposed region of visual cortex. After AAV injection, a glass window was used to cover the exposure; surgical-grade cyanoacrylate glue (VetBond) was used to seal the exposure and fix the glass window in place. Animals were allowed to recover for at least 3 weeks before imaging experiments were performed.

### In-vivo Calcium imaging

Calcium imaging was performed under isoflurane anesthesia. After anesthesia induction, the animal was head-fixed in a custom-built imaging stage and rested in a prone position on a fabric bed. A far infrared warming pad with closed-loop feedback control (RT-0515, Kent Scientific) was applied to maintain the animals’ body temperature during the whole experiment. The animal was anesthetized under 1% isoflurane during the whole imaging and stimulation period, with a maximum period of 1 hour.

An upper-right epi-fluorescence imaging system was used to image the calcium signal in the mouse cortex. The imaging system is modified from an Olympus BX51WI microscope. An excitation LED is used for calcium imaging (CoolLED pE-300), and the fluorescence signal is collected by a CMOS camera (C11440-36U, Hamamatsu ORCA-spark Digital CMOS Camera). The imaging frame rate is 20 Hz.

### Visual stimulation

A cranial window was installed on the mouse visual cortex through the procedure mentioned previously in the “Cranial window implantation and virus injection” section. The mouse was head fixed under the microscope and was anesthetized under 1% isoflurane during the whole imaging and stimulation period, with a maximum period of 1 hour. The brain activities in the visual cortex were recorded by the microscope through the procedure mentioned in the “in vivo calcium imaging” section. A far infrared warming pad with closed-loop feedback control (RT-0515, Kent Scientific) was applied to maintain the animals’ body temperature during the whole experiment. A white LED (MWWHL4, Thorlabs) was coupled into a 400 μm multimode fiber (FT400EMT, Thorlabs) for visual stimulation. The fiber was pointed to the mouse eye, and the distance between the fiber tip and the mouse’s eye is less than 5 mm. The LED was set to send 6 pulses in 2 s, with each pulse lasting 100 ms, and started to blink 5 s after the camera started recording. The resulting brain activities in the visual cortex were then recorded by the fluorescence camera. The stimulation was repeated 10 times for each animal. Three animals were tested in total.

### Blood-mediated optoacoustic stimulation

The animal was prepared in the same way as the “visual stimulation” section. To perform BOAS, the mouse was head fixed under the microscope and was anesthetized under 1% isoflurane during the whole imaging and stimulation period, with a maximum period of 1 hour. The brain activities in the visual cortex were recorded by the microscope. A far infrared warming pad with closed-loop feedback control (RT-0515, Kent Scientific) was applied to maintain the animals’ body temperature during the whole experiment. A 532 nm, 0.5 ns pulsed laser was weakly focused onto the mouse visual cortex with a diameter of ∼375 μm. The repetition rate of the laser was set to 20 kHz. Laser power and duration were determined by the different needs of experiments, which are mentioned in the results section. The laser was turned on for stimulation 5 s after the camera started to record. The resulting brain activities in the visual cortex were then recorded by the fluorescence camera. The stimulation was repeated 10 times for each animal. Three animals were tested in total.

### Histology examination

After the stimulation session, the mouse was sacrificed immediately, and then was perfused transcardially with phosphate-buffered saline (PBS, 1X, PH 7.4, Thermo Fisher Scientific Inc.) solution and 10% formalin. After fixation, the brain was extracted and fixed in 10% formalin solution for 24 h. The fixed mouse brain was immersed in 1X PBS solution. The brain was embedded in paraffin and sliced at a thickness of 5 µm to obtain coronal sections. Slicing and standard H&E staining were performed at Mass General Brigham Histopathology Research Core. Histology images were acquired with a VS120 Automated Slide Scanner (Olympus).

## Author Contributions

GC, J-XC and CY: drafting and refining the manuscript. GC, ML: built the imaging platform for calcium imaging and performed PA stimulation experiments. MT, KK: advised on surgical preparation of the animals and calcium imaging. KK, ML, XG, CM, NZ: preparing cranial windows on animals. DS, YL, HZ: help with the experiments. FC: help with animal breeding.

## Acknowledgement

This work was supported by the National Institute of Health, United States, through the BRAIN Initiative R01 NS109794 to J-XC and CY, BRAIN/NEI R21 EY035437 and NEI1R21EY036579 to CY.

## Data Availability

The data that support the findings of this study are available from the corresponding author upon reasonable request.

