## Supplementary Information for "Blood-Mediated Opto-Acoustic Stimulation of Brain Cortex at Sub-millimeter Precision"

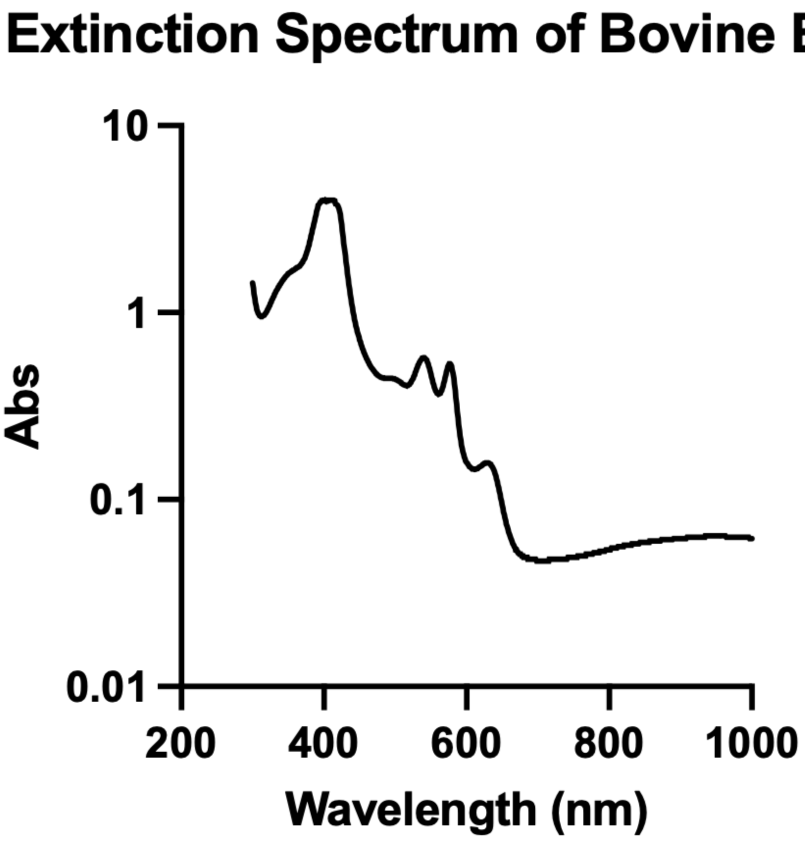


**Supplementary Figure S1.** Extinction spectrum of bovine blood measured by a spectrometer. Abs: absorbance.

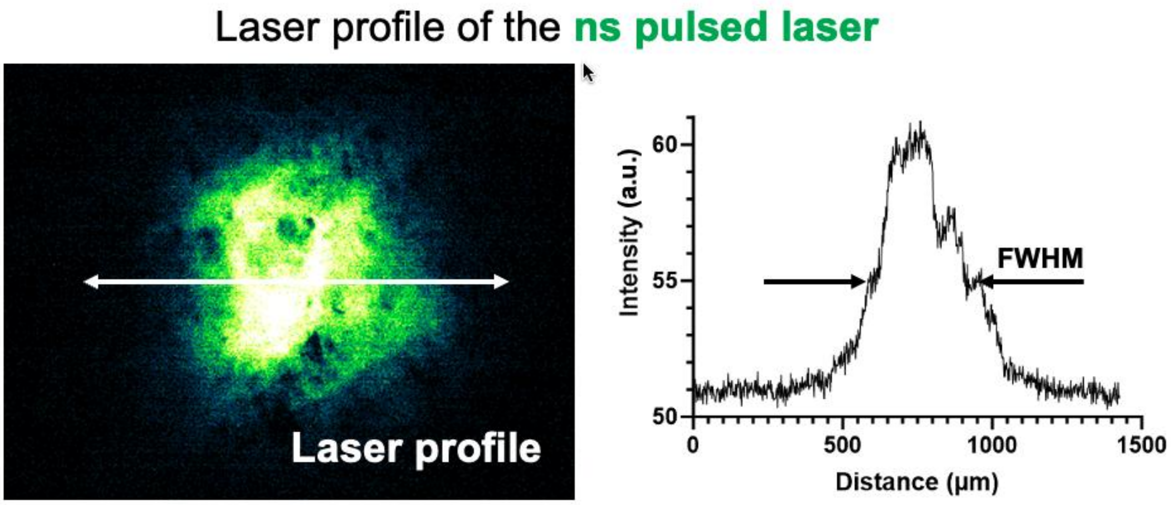


**Supplementary Figure S2. Intensity profile of the ns pulsed laser.** The FWHM (Full Width of Half Maximum) of the laser spot is ~375  μm.

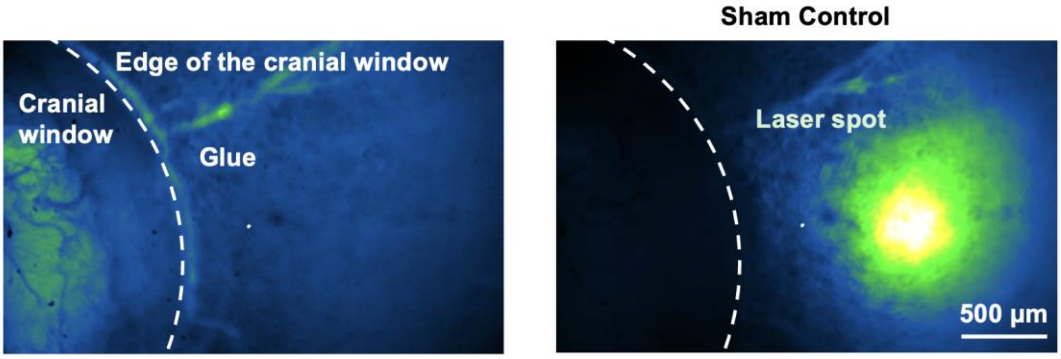


**Supplementary Figure S3.** The laser spot is moved out of the imaging area to perform the sham control.

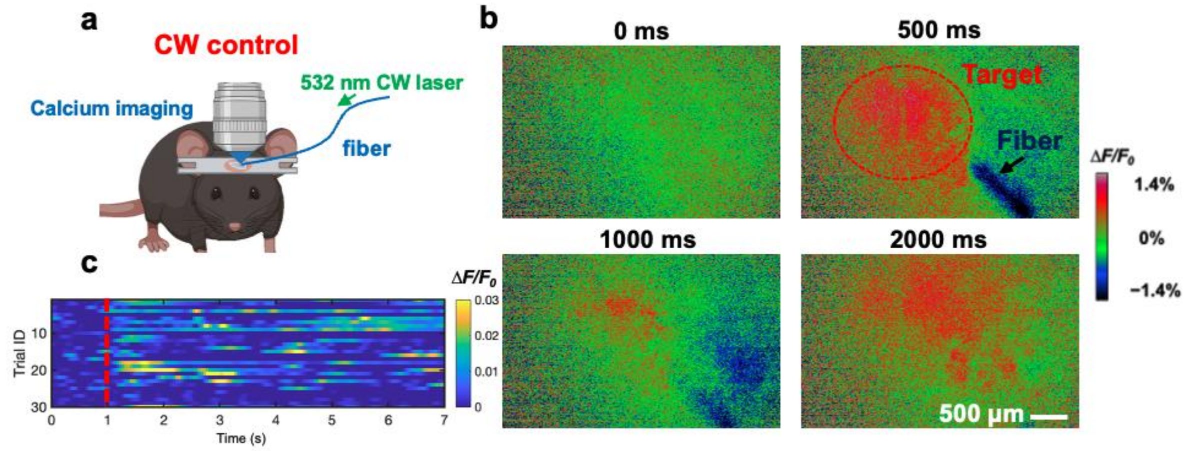


**Supplementary Figure S4. Photothermal stimulation of the brain using a 532 nm CW laser. a**. Schematic of photothermal stimulation using a multimode fiber and a 532 nm CW laser. Power on the sample: 200 mW. Laser duration: 100 ms **b**. Representative calcium imaging results of the photothermal stimulation on the visual cortex. **c**. Trial-by-trial traces of photothermal stimuli.
